# Reverse-engineering β-Arrestin Bias in the δ-Opioid Receptor

**DOI:** 10.1101/2025.11.27.690850

**Authors:** Tomasz M. Stepniewski, Manel Zeghal, Istvan Szabo, Maria Marti-Solano, Ismael Rodriguez-Espigares, Elizaveta Korchevaya, Mariona Torrens-Fontanals, Gianni de Fabritiis, Slawomir Filipek, Patrick M. Giguère, Jana Selent

## Abstract

G protein–coupled receptors (GPCRs) form the largest family of cell-surface receptors and remain prime targets in drug discovery. A central challenge in modern GPCR drug discovery is understanding and exploiting biased agonism: the ability of ligands to favor signaling via therapeutically beneficial pathways while avoiding those that trigger side effects. Therefore, pinpointing the structural determinants of signaling bias is crucial for rational drug design. Biased agonists are particularly compelling for targeting opioid receptors, as in this family, ligands that limit β-arrestin (β-arr) recruitment are believed to preserve analgesia while reducing respiratory depression and addiction liabilities.

Here, we use extensive all-atom molecular dynamics (MD) simulations to dissect signaling bias in the δ-opioid receptor (δOR). Focusing on a receptor mutant with a strong β-arr bias, we employed a reverse-engineering approach to reveal the conformational mechanisms that promote β-arr recruitment. Building on these insights, engineer new mutations that reshape the receptor’s signaling profile. Importantly, this approach allowed us to pinpoint signaling bias to motions of a single microswitch and identify how structural receptor motions induced by the mutations and ligand contacts cooperate to promote a specific functional response. In this proof-of-concept study, we not only provide structural insights into δOR pharmacology but also demonstrate how computational methods can be leveraged to probe structural mechanisms of signaling specificity across GPCRs, paving the way for the rational design of tailored receptor variants and novel, safer, and more effective therapeutics.

## Introduction

G protein-coupled receptors (GPCRs) constitute the largest family of cell surface receptors in the human genome(*1*), ubiquitously expressed across nearly all human cell types and integral to numerous physiological processes. Their pharmacological relevance is underscored by the fact that more than one-third of all marketed drugs target this protein family(*2*). A major challenge in GPCR pharmacology is the development of ligands with functional selectivity, or signaling bias, which can preferentially engage therapeutic pathways while avoiding those linked to side effects. Such biased agonists hold promise as safer and more efficacious therapeutics(*3*). However, the rational design of biased ligands remains elusive, as signaling bias arises from subtle perturbations within an extensive intramolecular allosteric network(*4–6*). Understanding mechanistically how local structural changes propagate through this network to reconfigure receptor conformations and effector coupling is therefore critical and remains a central challenge in GPCR drug discovery.

To address this question, we introduce a reverse-engineering approach to gain a deeper understanding of GPCR signaling. We take advantage of a rare type of mutation—a single amino acid substitution that rewires signaling in the δ-opioid receptor (δOR) and transforms a series of antagonists into potent β-arrestin (β-arr)-biased agonists(*7*). Rather than only describing bias as an emergent property, our goal was to deconstruct how specific structural perturbations reshape the receptor’s conformational and signaling landscape and then engineer these changes to evaluate their impact on the functional response. Based on previous experiences using *in silico* methods to understand GPCR signaling(*8–10*), we extensively sample this system through molecular dynamics (MD) simulations, mapping the receptor’s intramolecular interaction network and identifying specific perturbations and microswitch rearrangements that drive signaling towards β-arrestin. Then, as a proof of concept, we use the gained insights to computationally design a novel set of mutations that engineer bias within the δOR, with some of the mutations fine-tuning the signaling response by perturbing a single microswitch.

All in all, by systematically engineering δOR variants with tunable β-arr bias, we establish a structural blueprint for controlling receptor signaling outcomes. Although the exact impact of β-arr recruitment on the pharmacological effects of opioid agonists is still under investigation (*11*, *12*), in the δOR(*13*) and the closely related prototypical receptor μOR, β-arr has been linked to drug side effects and tolerance. As the δOR is an attractive target for pain management(*14*) as well as anxiety and mood disorders(*15*, *16*), our findings provide potential insights for the rational design of safer, pathway-selective opioid therapeutics.

## Results and discussion

### Flexibility analysis of MD simulations provides mechanistic insight into β-arrestin bias in the δOR

To reverse engineer the mechanics of signaling in the δOR, we focused on the D2.50A δOR. This mutation converts a series of δOR antagonists (naltrindole among them) into potent β-arr-biased agonists(*7*). Such a functional outcome is surprising, as, despite some reports(*17–19*), a single-point mutation inducing an antagonist-agonist shift is uncommon. D2.50 is a conserved residue in Class A GPCRs and a principal component of the sodium ion binding site. Ions binding within this region are known to allosterically modulate class A GPCRs and stabilize inactive-like conformations(*20*, *21*). To investigate the mechanistic basis of the effect of this mutation, we performed extensive molecular dynamics simulations (100 replicates × 128 ns) of both the wild-type (WT) and D2.50A δOR in complex with naltrindole. In line with the experimental results, we included an allosterically bound sodium only in the WT system, as the D2.50A δOR is sodium-insensitive(*7*). Initially, we wanted to identify disturbances in macrostructural elements; we studied flexibility using backbone root-mean-square fluctuation (RMSF). This analysis revealed several regions with altered dynamic behavior between the two systems (Fig. 1A). Notably, the intracellular segments of TM5 and TM6 exhibited reduced fluctuations in the D2.50A mutant, while the extracellular regions of TM1 and TM5 showed increased flexibility. Interestingly, our analysis also identified enhanced fluctuations in the transmembrane domain in an extended (spanning approximately two helical turns) intracellular part of TM7. The magnitude of the fluctuations in TM7 exceeded that observed at the site of the D2.50A mutation itself, which typically undergoes the greatest perturbation during the initial equilibration phase. Increased mobility of structural elements is linked to agonist activity(*22*); thus, we speculated that the observed gain in mobility could be linked to the increase in β-arr recruitment. To better understand how the D2.50A mutation induces this structural instability, we examined specific residues in the TM7 segment. This region contains multiple polar and highly conserved(*23*) residues (Fig. 1B), which can form interactions with neighboring side-chains. Analysis of polar contacts within this segment (Fig. 1C) highlights that the D2.50A mutation disrupts two direct polar interactions between TM2 and TM7 (D2.50 and S7.46, as well as D2.50 and 7.49). Furthermore, mapping water occupancy in the WT system (Fig. 1D, left) revealed the presence of an internal water network centered around the sodium ion. This network mediates multiple interactions between TM2 and TM7 (i.e., N7.45 and D2.50) or between TM2 and TM3 (S3.39 and D2.50). Unsurprisingly, the D2.50A mutant, which lacks the water-coordinating ion, exhibits a destabilized water network (Fig. 1D, right), leading to the disruption of multiple inter-helical contacts. This is evidenced by the loss of four stable waters when comparing the WT and D2.50A water networks. Overall, our analysis revealed that within the δOR (Fig. 1E), D2.50 engages in several direct and water-mediated inter-helical interactions (Fig. 1F), primarily connecting TM2 and TM7. We therefore speculate that the loss of these interactions underlies the destabilization of TM7 (Fig. 1G–H), potentially contributing to the observed β-arr bias in the studied systems. In such a scenario, TM7 stability could act as a tunable functional switch enabling the engineering of β-arr bias within the δOR.

**Figure 1.**
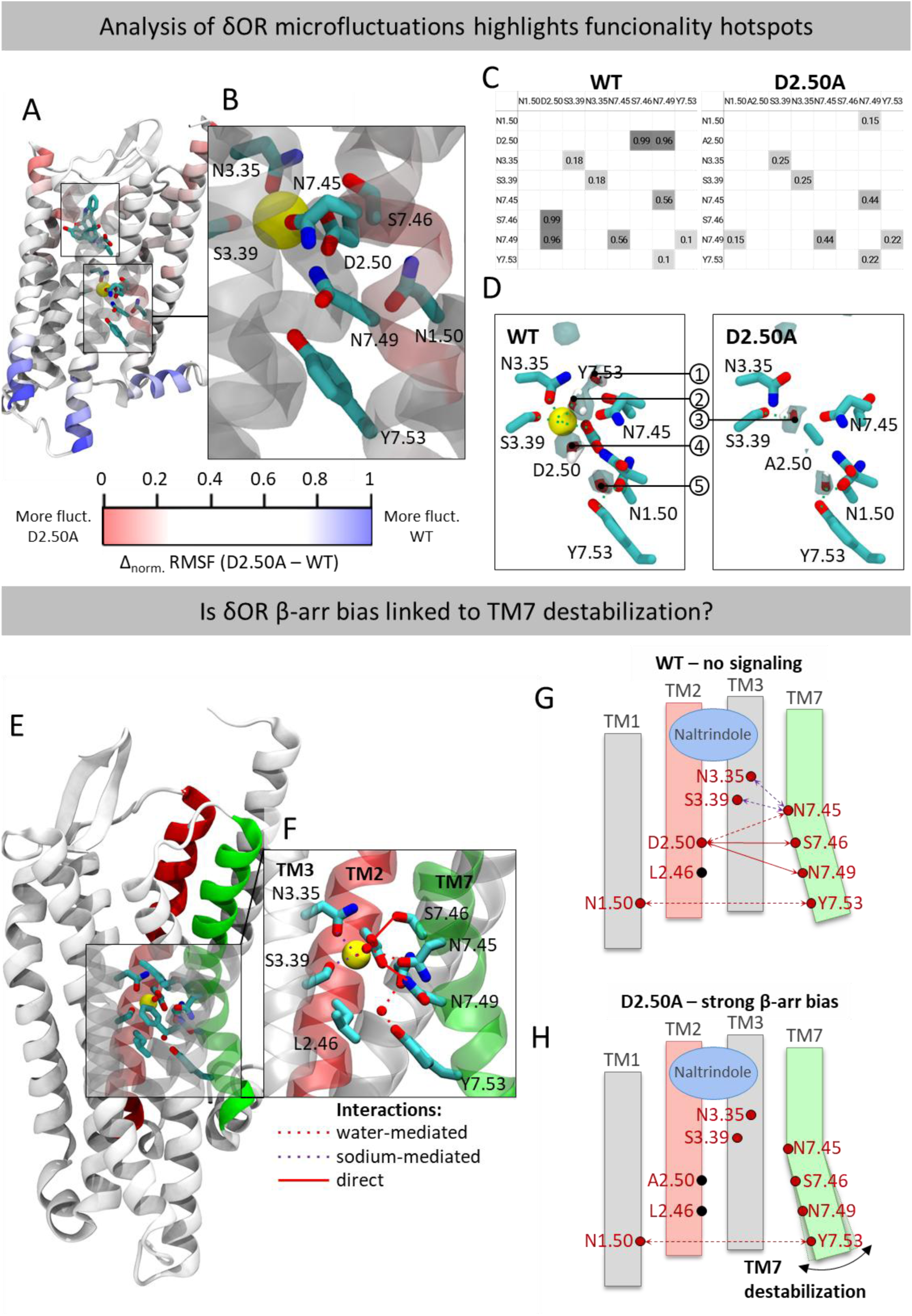
Putative macro-structural determinants of β-arrestin bias in the δOR. **(A)** Backbone flexibility was compared between simulations (100 × 128 ns) of the WT and D2.50A δOR bound to naltrindole. Flexibility was quantified as the Root Mean Square Fluctuation (RMSF) of Cα atoms, and residue-wise ΔRMSF values were obtained by subtracting WT values from those of D2.50A. The resulting values were normalized by feature scaling (see Methods). The structural model is colored such that regions more flexible in WT are shown in blue, and those more flexible in the β-arrestin–biased D2.50A variant in red. **(B)** Structural view of the conserved 2.50 position in WT δOR. Residues are shown as licorice, with the bound sodium ion depicted as a yellow sphere. **(C)** Comparison of direct polar contacts in residues neighboring the conserved 2.50 position between WT and D2.50A. Interactions were identified using the get-contacts package. **(D)** Water networks in WT and D2.50A systems. Water occupancy was approximated from volumetric maps of water oxygen atoms, depicted at an occupancy threshold of 0.15. **(E)** Schematic representation of the polar network connecting TM2 and TM7 in δOR, with a detailed view **(F)** in the vicinity of residue D2.50. Schematic comparisons of this polar network are shown for WT **(G)** and D2.50A δOR **(H),** differentiating direct polar interactions (red lines) and those mediated by water (red dotted lines) or sodium (purple dotted lines)

### Reverse-engineering β-arrestin bias in the δOR on the macrostructural level

To verify the mechanistic hypothesis that TM7 mobility drives the β-arr bias in the δOR and that this bias can be engineered by targeted TM7 destabilization, we designed mutations predicted to induce this structural change. The first group of mutations was tailored to promote TM7 flexibility by disrupting TM2-TM7 interactions (N7.45A, S7.46A, and N7.49A), as guided by the identified inter-helical interaction networks (Fig. 1E). The P7.50A mutant was also designed to destabilize TM7, in this case by inducing changes in the receptor backbone rather than directly changing inter-helical interaction patterns. Lastly, we designed one control mutant: L2.46A. Although the 2.46 position is near TM7, it does not interact directly with the TM7 helix. Thus, we speculated that a L2.46A mutation would not change inter-helical interaction patterns and TM7 stability and therefore not impact β-arr recruitment.

The structural impact of each mutant was computationally evaluated using MD simulations (50 replicates × 128 ns). Both visual inspection (Fig. 2A) and RMSF analysis (Fig. 2B) confirmed the ability of the 4 designed mutants, TM7 (N7.45A, S7.46A, N7.49A, and P7.50A) to destabilize TM7. Also, as predicted, the designed control mutant (L2.46A) did not affect TM7 flexibility. Having structurally characterized the designed variants, we next evaluated their impact on the signaling response to naltrindole. The Gαi activation assay confirmed that none of the studied mutants promoted G-protein signaling (Fig. 2C). In contrast, and consistent with our initial structural hypothesis, each of the TM7-destabilizing mutants enhanced β-arr recruitment to varying degrees, with the P7.50A mutant producing the most pronounced effect (Fig. 2D). Further in line with our hypothesis, the L2.46A mutant, which does not perturb TM7 stability in our simulations, showed no influence on β-arr coupling, not changing the profile of naltrindole form an antagonist. These results demonstrate that by rationally engineering TM7 destabilization (similar to the D2.50A mutation), we can selectively bias the naltindole/δOR complex towards β-arr. Thus, this reverse-engineering study highlights TM7 as a critical hotspot for β-arr bias and establishes a direct causal link between local helix destabilization and altered signaling outcomes.

**Figure 2.**
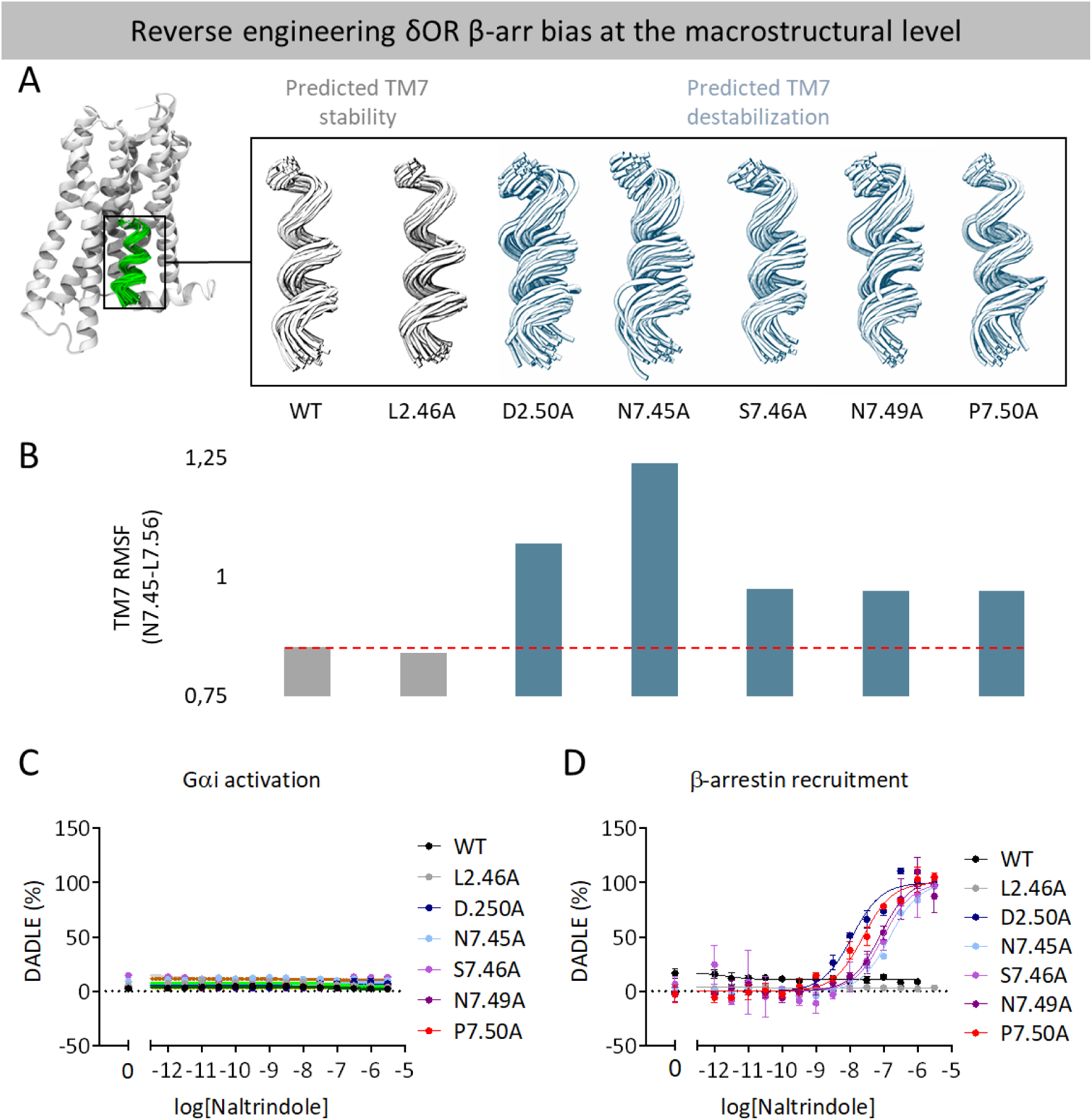
TM7 instability induces β-arrestin bias in the δOR/naltrindole complex. **(A)** Structural snapshots of TM7 backbone conformations (residues 7.45 to 7.56) explored during 100 x 128 ns (WT and D2.50A) and 50 x 128 ns (other mutants) of simulation time (one snapshot every 40ns). Systems with reduced TM7 flexibility are colored grey. **(B)** RMSF values for the TM7 backbone (residues 7.45 to 7.56) were calculated for all simulated systems. Systems with reduced TM7 flexibility are colored grey. **(C)** Normalized concentration-responses of δ-OR-mediated Gα_i_ signaling induced by naltrindole **(D)** and normalized concentration-responses of δ-OR-mediated β-arrestin recruitment induced by naltrindole. Results were quantified as described in the methods section.

Although, to our knowledge, no MD project has studied the mutation-induced transition of an antagonist into an agonist, studies in other class A and B GPCRs show similar trends for different receptors. NMR studies of the prototypical β2-adrenergic receptor reveal a similar trend, indicating that β-arr-biased agonists primarily perturb TM7, while agonists promoting G protein bias primarily modulate TM6(*24*, *25*). A similar trend has been observed in fluorescence experiments using the vasopressin 2 receptor(*26*). Finally, the involvement of TM7 in β-arr bias in GPCRs is supported by crystallographic data on the serotonergic receptors. In the structure of the 5-HT2B receptor bound to ergotamine, where it acts as a β-arr biased agonist, an activation-related shift of the intracellular segment of TM7 can be observed, which is more pronounced than the one found in the 5-HT1B receptor structure, where ergotamine acts as a balanced agonist(*27*). Within opioid receptors, TM7 has been linked to β-arrestin recruitment in both the μOR and κOR. For μOR, a study comparing G protein-biased and balanced agonists using a combination of MD simulations and NMR, found links between the conformation of TM7, ICL1, and helix 8 and arrestin recruitment(*28*). Furthermore, pharmacological analysis of the same receptor reveals that the engagement pattern of the extracellular part of TM7 is a strong predictor of β-arr recruitment(*29*). For the κOR, computational results suggest that an alternative confirmation of TM7 drives the β-arr bias of the agonist WMS-X600(*30*). Considering this, our results, which link TM7 to β-arr bias and recruitment in δOR signaling, are supported by data on other GPCRs and suggest a general mechanism, conserved in class A GPCRs, that could be exploited in the search for new biased agonist therapies. Importantly, uncovering this mechanism in the δOR highlights the power of our reverse-engineering approach to reveal previously unrecognized determinants of functional selectivity.

### Reverse-engineering β-arr bias in the δOR at the microswitch level

Although our results demonstrate a link between β-arr bias and TM7 destabilization, we wanted to evaluate whether we could pinpoint structural mechanisms of bias at a lower structural resolution. To dissect the impact of TM7 on a microswitch level, we focused on specific residues within the helix. TM7 contains several conserved microswitches, most notably Y7.53, a critical component of the NPxxY motif that plays a central role in GPCR activation(*31–34*). Comparing the WT and D2.50A systems, we observed a marked increase in the conformational flexibility of Y7.53 in the D2.50A receptor (Fig. 3A), consistent with a local destabilization of the TM7–helix interface and altered water-mediated interactions. Specifically, while in the WT system Y7.53 adopts primarily a conformation in which the hydroxyl group is oriented towards the internal receptor space (conf 1), in the D2.50A system the residue also assumes a secondary conformation, where the hydroxyl group of Y7.53 is rotated toward the intracellular space (conf 2) (Fig. 3B, top vs middle). Interestingly, among the available X-ray and Cryo-EM structures of the δOR, two have been crystallized, bearing mutations that strongly promote β-arr recruitment and only moderately promote G-protein recruitment(*35*). Thus, the crystallographic data effectively capture a static image of δOR in a β-arr-biased state (Fig. 3C), potentially allowing us to pinpoint specific structural re-arrangements specifically linked to β-arr bias. In both structures, Y7.53 adopts a conformation where the hydroxyl group is oriented towards the intracellular space, similar to conformation 2 (Fig. 2B, middle vs bottom). Driven by this discovery, we hypothesized that TM7 flexibility in δOR can promote β-arr bias by altering the conformational ensemble of Y7.53. For additional verification, we analyzed Y7.53 mobility across other simulated systems. To differentiate between conformation 1 and conformation 2, we used the distance between the oxygen of the polar Y7.53 OH group and the Cα atom of N8.49 (Fig. 3D), quantifying as conformation 2 orientation of Y7.53 where the distance is lower than 10Å (Fig. 3E). Consistent with our predictions, we find that in systems without β-arr bias (WT and L2.46A), Y7.53 remained almost exclusively in conformation 1. In contrast, in systems exhibiting β-arr bias, Y7.53 more frequently assumes conformation 2 (Fig. 3F). Motivated by these findings, we aimed to test whether β-arr bias could be engineered at the microswitch level by directly engineering a shift of Y7.53 towards conformation 2. In the inactive δOR structure (PDB: 4N6H), Y7.53 is primarily stabilized through water-mediated contacts with N1.50 and hydrophobic contacts with V6.40 (Fig. 3G). To induce Y7.53 destabilization, we virtually mutated both N1.50 and V6.40 into alanine and monitored the Y7.53 conformation. While N1.50A promoted TM7 flexibility, V6.40 did not significantly alter the flexibility of the segment (Fig. 3H). Importantly, both mutations shifted the conformational equilibrium of Y7.53 towards conformation 2 (Fig. 3I), with V6.40A inducing a markedly stronger shift than N1.50A (2,13% vs 5,6%). In line with the simulation-derived predictions, experiments confirmed that both mutations induce β-arr bias, with V6.40A inducing the strongest bias among all studied mutants, even considering the previously identified D2.50A(*7*) (Fig. 3J-K). These results further support the role of the Y7.53 conformation, particularly conformation 2, in driving β-arr bias in δOR, highlighting the residue as a signaling microswitch. Furthermore, they demonstrate how, by profoundly analyzing macro-structural changes in a receptor, it is possible to pinpoint micro-structural switches and engineer bias by shifting the conformational equilibria of a single residue.

**Figure 3.**
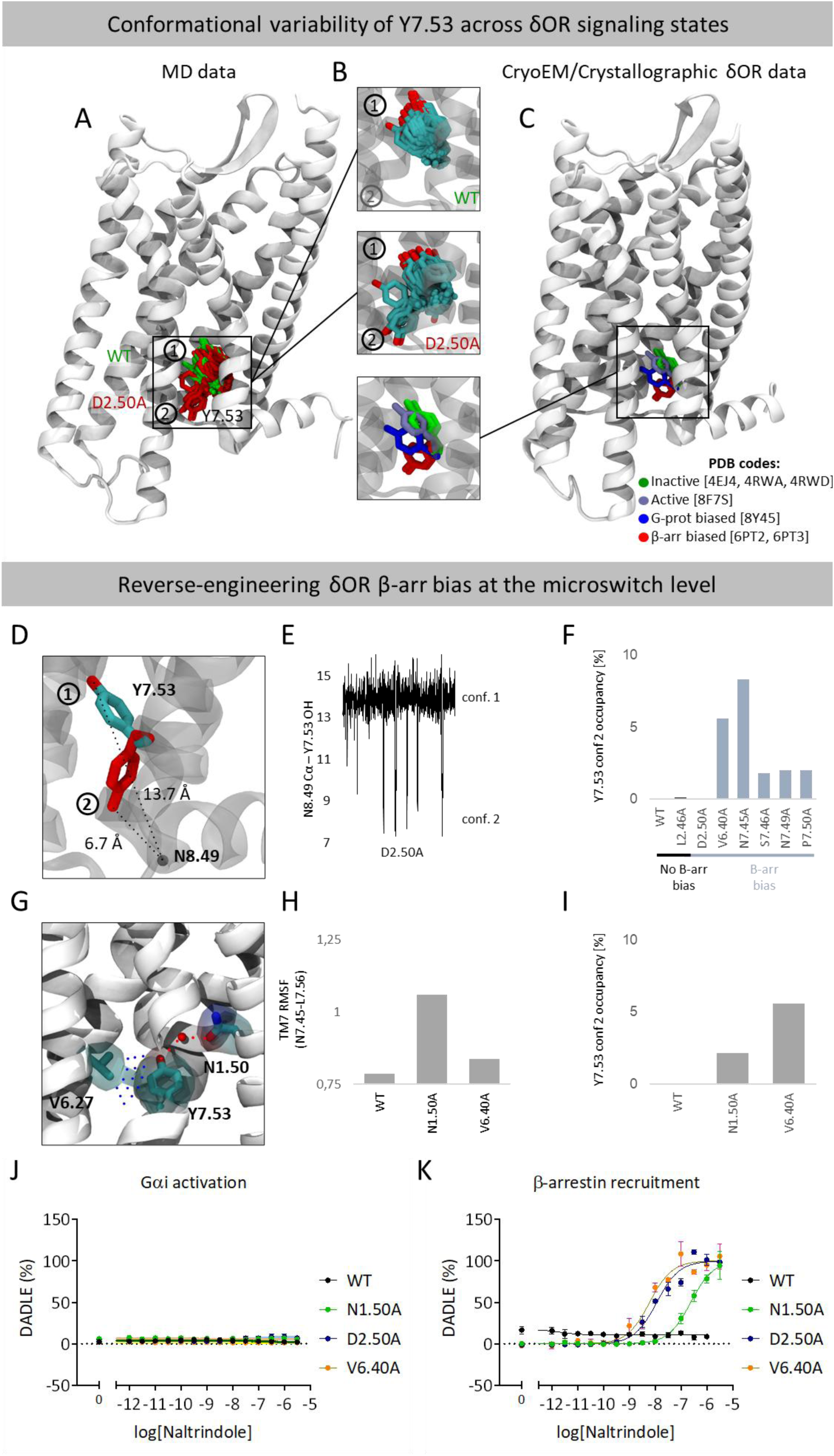
Impact of the Y7.53 on β-arrestin bias in the δOR – identification of a functional selectivity microswitch. **(A)** Conformations of Y7.53 observed in WT (green) and D2.50A (red) simulations (one snapshot every 40ns). Based on the orientation of the residue, conformations have been classified into conformation 1 (oriented towards the orthosteric binding site) and conformation 2 (oriented towards the intracellular site). **(B)** Conformations of Y7.53 observed in MD simulations of the WT δOR (top), D2.50A (middle), and X-ray and CryoEM structures (bottom). Conformations are colored according to the functional state of the receptor **(C)** with structures in an inactive state being colored green (PDB codes: 4EJ4, 4RWA, 4RWD), in an active state colored purple (PDB code: 8F7S), in a G protein biased state colored blue (PDB code: 8Y45) and β-arr biased state colored red (PDB codes: 6PT2 and 6PT3). **(D)** Differentiating conformation 1 and conformation 2 based on the distance between the OH atom of Y7.53 and Cα of N8.49 **(E)** Conformations of Y7.53 explored in the D2.50A system, based on the Y7.53 OH and Cα of N8.49 distance **(F)** Percentage of frames where Y7.53 assumes conformation 2 (based on the Y7.53 OH and Cα of N8.49 distance) in unbiased (black) and β-arr biased (blue) systems. **(G)** Residues stabilizing the Y7.53 conformation in the δOR with polar (red) and hydrophobic (blue) interactions. **(H)** RMSF values for TM7 backbone (residues 7.45 to 7.56) calculated for N1.50A, V6.40A δOR (50 x 128ns) and WT (100 x 128ns(. **(I)** Percentage of frames where Y7.53 assumes conformation 2 in WT, N1.50A and V6.40A δOR. **(J)** Normalized concentration-responses of δ-OR-mediated Gα_i_ signaling induced by naltrindole **(K)** and normalized concentration-responses of δ-OR-mediated β-arrestin recruitment. Results were quantified as described in Methods section.

Notably, the involvement of the Y7.53/NPxxY motif in β-arr bias has been studied in other receptors. Computational studies suggested that in the angiotensin (AT1R), μOR, and cannabinoid 1 (CB1R) receptors, the conformation of the NPxxY is linked to β-arr bias (*36–38*). The “β-arr-biased” conformation can be characterized by a strong inward movement (μOR and AT1R) or rotation (CB1 and AT1R) of the intracellular segment of TM7. Additionally, in line with our findings in the AT1R, the proposed β-arr biased conformation demonstrates an intracellular rotation of both Y7.53 and R3.50. Thus, alternative studies on different receptors suggest that Y7.53 is involved in β-arr bias, supporting the validity of our findings.

### Engineering β-arr bias in the δOR requires cooperative TM6 stabilization and TM7/Y7.53 flexibility

Although the mechanistic link between TM7 flexibility and β-arr bias provides insight into signaling within the δOR, it does not explain why in all studied systems β-arr recruitment is antagonist dependent. As highlighted by previous experimental data (*26*), the D2.50A mutation in the δOR on its own induces only a very modest β-arr bias, and robust recruitment of β-arr requires binding of naltrindole or other antagonists. Thus, to verify how the presence of naltrindole shifts this signaling response towards β-arr, we simulated the D2.50A δOR in its apo-state (100 replicates x 128 ns).

Comparison of backbone RMSF values between the apo and naltrindole-bound D2.50A δOR revealed reduced flexibility in the antagonist-bound receptor (Fig. 4A). The most pronounced changes occurred in the extracellular end of TM6 and the intracellular ends of TM5 and TM6. Interestingly, the altered flexibility pattern of TM6, both in the orthosteric binding site (the naltrindole binding site) and the intracellular region, would suggest that the change in flexibility occurs due to TM6/naltrindole contacts, which are propagated across TM6. These results are in line with general knowledge about the pharmacology of class A GPCRs, where agonists typically engage TM5 and TM6 within the orthosteric site, inducing intracellular rearrangements—most notably the outward movement of TM6—that create space for G protein binding(*31*, *39–41*). Conversely, antagonists such as naltrindole bind to inactive-like conformations where the intracellular part of TM6 remains closer to the helical bundle, preventing G protein coupling. Thus, the ability of naltrindole to stabilize both TM5 and TM6 is in line with its functional profile as an antagonist.

**Figure 4.**
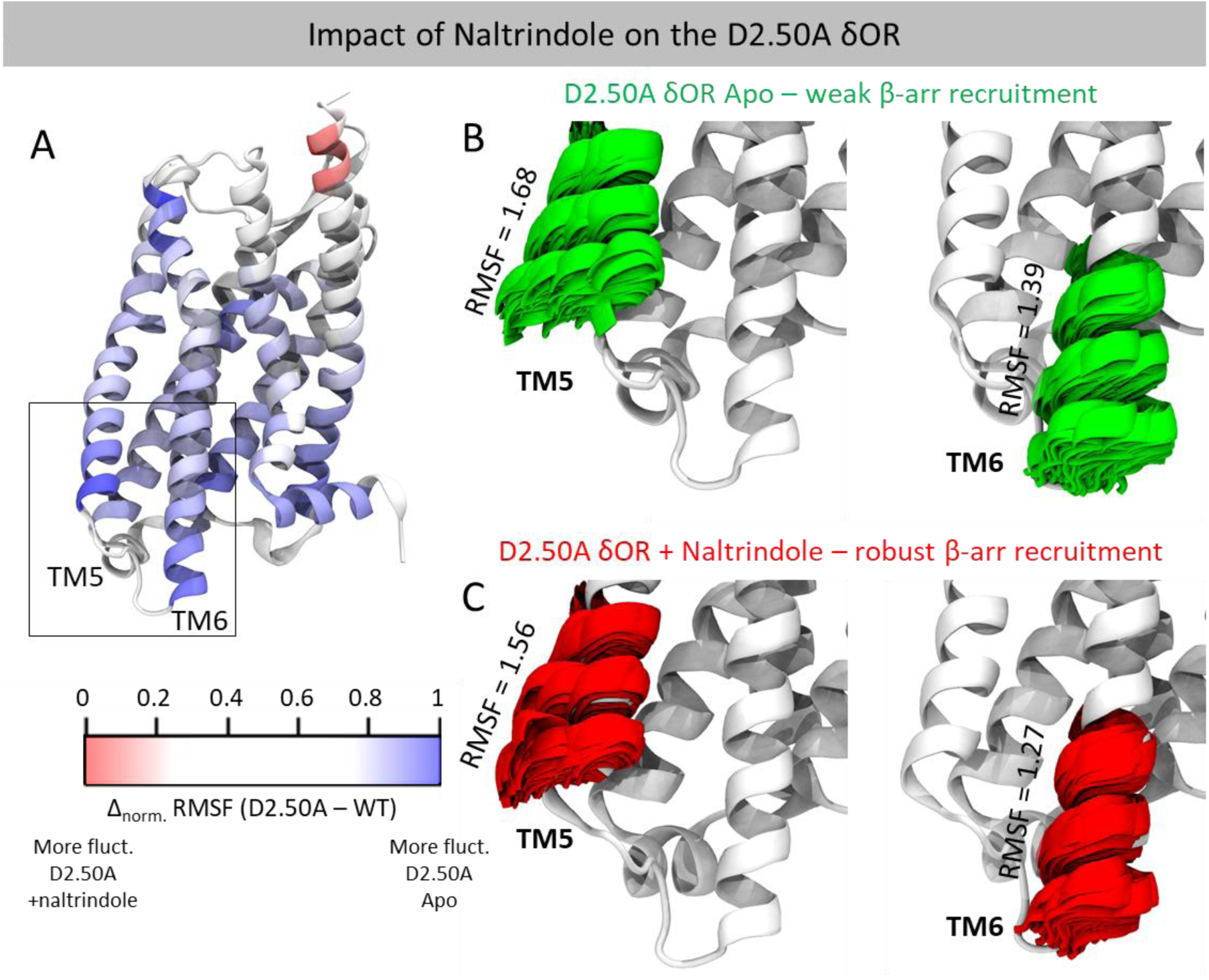
Structural comparison of naltrindole-bound and apo δOR. **(A)** Backbone flexibility was compared between simulations (100 × 128 ns) of the naltrindole-bound D2.50A and Apo δOR. Flexibility was quantified as RMSF of Cα atoms, and residue-wise ΔRMSF values were obtained by subtracting naltrindole-bound system values from those of Apo system. The resulting values were normalized by feature scaling (see Methods). The structural model is colored such that regions more flexible in WT are shown in blue, and those more flexible in the β-arrestin–biased D2.50A variant in red. **(B)** Structural snapshots of TM5 and TM6 flexibility in the Apo D2.50A δOR with associated RMSF values of Cα atoms (K6.24-T6.34, one snapshot every 64ns). **(C)** Structural snapshots of TM5 and TM6 flexibility in the naltrindole-bound D2.50A δOR with associated RMSF values of Cα atoms (K6.24-T6.34, one snapshot every 64ns).

As shown earlier (Fig. 3G), alterations in the TM6–TM7 interaction network can shift the conformational equilibrium of Y7.53. Motivated by this observation, we examined the dynamics of Y7.53 (Fig. 3D). Consistent with our previous data, in the apo D2.50A receptor (which does not exhibit β-arr bias), Y7.53 shows a markedly reduced propensity to adopt conformation 2 compared to the naltrindole-bound D2.50A δOR (Fig. 5A-B). To determine whether this altered mobility of Y7.53 is linked to a specific TM6 conformation, we analyzed TM6 in frames where Y7.53 adopts conformation 2. In this subset, TM6 displays a pronounced shift from TM7 (Fig. 5C) where Y7.53 is located. This shift becomes more evident when comparing the selected frames with the distribution observed in the naltrindole-bound D2.50A δOR (Fig. 5D). Consistently, comparison of apo and naltrindole-bound conformations reveals that naltrindole induces an outward shift of the intracellular segment of TM6 (Fig. 5E-F). Taking this into account, it is tempting to speculate that naltrindole is crucial for the studied mutants to elicit β-arr bias in the δOR, as it stabilizes a specific conformation of TM6, which allows a downward orientation of Y7.53 (Fig. 5G-H). To further support the notion that a specific positioning of TM6 is required to promote signaling bias, we examined the effect of the mutation that most strongly enhances β-arr recruitment (V6.40A) on the signaling profile of the extensively studied δOR agonist DADLE(*42*). As DADLE induces δOR activation, it likely stabilizes a distinct conformation of the intracellular coupling site compared to the antagonist naltrindole, particularly involving the intracellular segments of TM5 and TM6. Interestingly, within the DADLE-bound system, the V6.40A mutation reduces G protein signaling. Furthermore, in contrast to naltrindole, this mutation does not promote β-arr recruitment in response to DADLE and, in fact, reduces it (Fig. 5I). Taken together, these observations suggest that engineering signaling bias within the δOR requires not only destabilization of TM7/Y7.53, but also conformational locking of TM6 in a position that increases the distance between TM6 and TM7. Such an arrangement of TM6 and TM7 enables Y7.53 to adopt conformation 2, which is associated with β-arr biased signaling. Interestingly, this hypothesis correlates with X-ray data, as in δOR structures captured in a β-arr biased conformation, TM6 is notably extended away from TM7 (Fig. S2). The proposed mechanism is also in line with data on the CB1 receptor. A study by Fay et al. showed that the activity of the positive allosteric modulator Org 27569, which induces β-arr bias, is linked to its ability to block TM6 mobility and potentiate the mobility of TM7(*43*). Furthermore, fluorescence spectroscopy revealed similar trends in the vasopressin V2 receptor, where agonists inducing G protein recruitment induced conformational shifts in TM6, whereas the β-arr-biased agonist SR121463 did not induce shifts in TM6 and only altered the conformation of TM7(*26*).

**Figure 5.**
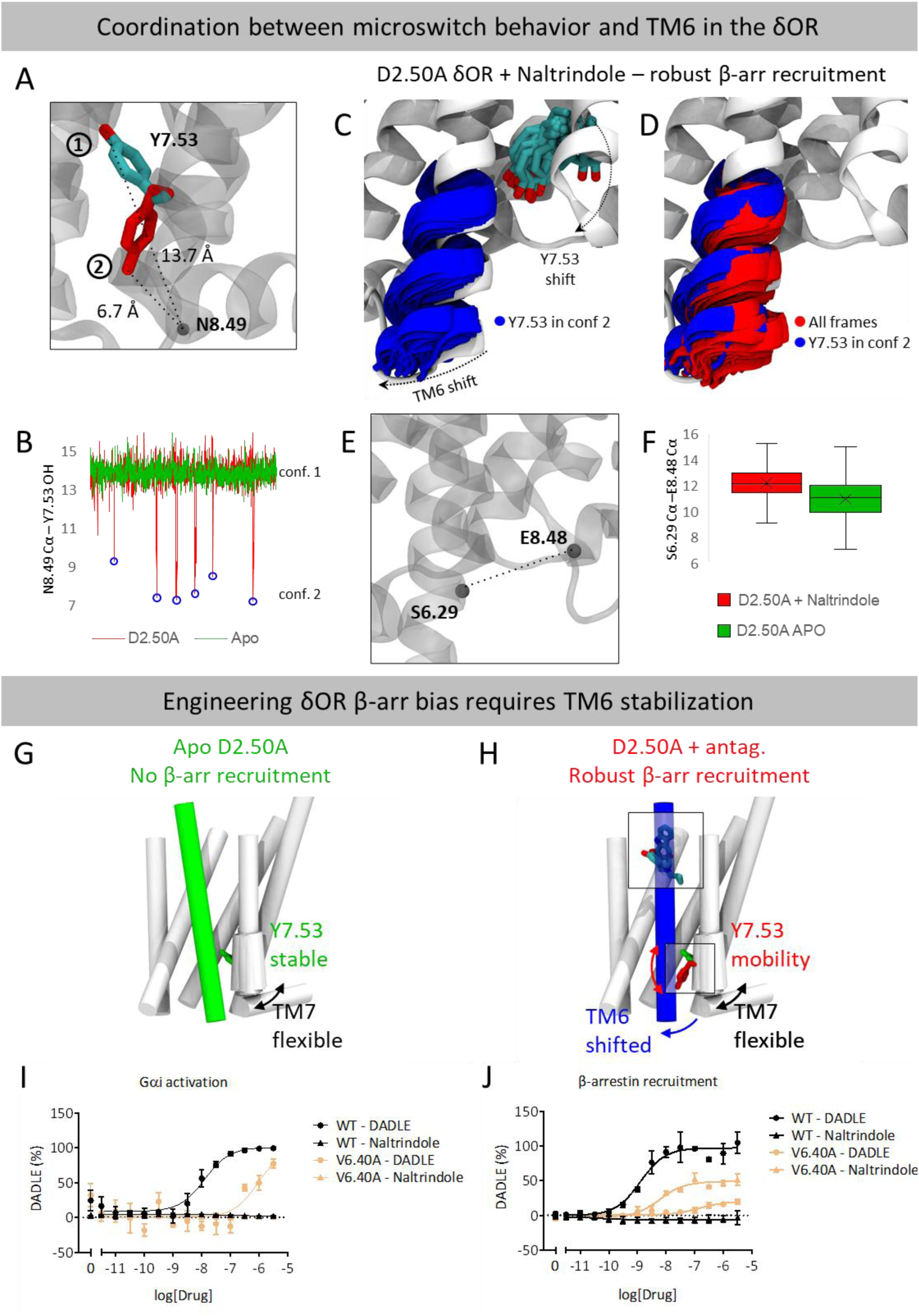
Structural insights into the impact of naltrindole on β-arrestin bias in the δOR. **(A)** Differentiating conf. 1 and conf. 2 based on the distance between the OH atom of Y7.53 and Cα of N8.49 **(B)** Conformations of Y7.53 explored in the D2.50A (red) and Apo (green) systems, based on the Y7.53 OH and Cα of N8.49 distance **(C)** Conformations explored by TM6 in the naltrindole-bound D2.50A δOR in frames where Y7.53 assume conf. 2 **(D)** Comparison of conformations explored by TM6 in the naltrindole-bound D2.50A δOR in frames where Y7.53 assume conf. 2 and in all frames (one snapshot every 64 ns) **(E)** Measuring the shift of TM7 using Cα of S6.29 and E8.48 **(F)** Comparison of TM6-shift between the naltrindole-bound and Apo D2.50A δOR **(H)** Schematic representation of the impact of naltrindole **(I)** Normalized concentration-responses of δOR-mediated Gα_i_ signaling induced by DADLE and naltrindole **(J)** and normalized concentration-responses of δOR mediated β-arrestin recruitment. Results were quantified as described in the Methods section.

## Conclusions

In this study, we aimed to reverse-engineer β-arrestin bias in the δ-opioid receptor by combining large-scale molecular dynamics simulations with rational mutagenesis. By first identifying macro-regions with altered flexibility and then focusing on the behavior of specific residues in the studied segment, we find that the D2.50A mutation destabilizes the sodium binding site, triggering conformational flexibility in TM7 and altering the ensemble of the conserved Y7.53 microswitch. By systematically targeting this region with designed mutations, we were able to reproduce and even slightly enhance β-arrestin recruitment, thereby demonstrating that signaling outcomes in GPCRs can be rewired through precise structural perturbations. Beyond characterizing a hotspot for functional selectivity, this reverse-engineering perspective highlights how structural elements collectively shape signaling outcomes (i.e., the interplay between TM6 and TM7) and demonstrates how an antagonist can be transformed into a potent biased agonist. While previous computational studies have provided fragmented insights into this receptor’s function(*44–47*), our work integrates multiple structural features into a coherent, high-resolution framework. Furthermore, we demonstrate how computational techniques can be utilized to understand and engineer signaling bias, facilitating the design of receptor variants with a defined signaling bias. In this way, receptor mutants like the newly identified V6.40A could serve as functional tools to experimentally explore the link between β-arr signaling and the unwanted side effects elicited by current drugs targeting the δOR.

From a pharmacological perspective, the obtained results provide novel structural guidance for designing functionally selective agonists targeting opioid receptors, particularly through the modulation of TM7 flexibility. Furthermore, recent drug discovery efforts on the OR have focused on developing bitopic ligands that can target both the orthosteric and the conserved sodium allosteric site(*48*, *49*). In this context, our study offers valuable mechanistic insights into how alterations in this region can fine-tune receptor signaling and would suggest that a ligand capable of stabilizing TM7 while allowing an intracellular conformational shift in the TM6 region could bias the receptor away from β-arr binding and, in this way, potentially display an improved therapeutic profile.

All in all, this study illustrates that receptor signaling bias can be reverse-engineered through targeted structural manipulations. By linking local flexibility changes to global signaling outcomes, we demonstrate that it is not only possible to understand, but also to redesign GPCR function at the molecular level. This approach transforms studying bias from a descriptive into a creative process—demonstrating that receptor activity can, quite literally, be reprogrammed by design.

## Methods

### Molecular dynamics simulations

All simulated systems were generated using the structure of the δOR bound to the antagonist naltrindole [PDB code: 4N6H](*7*). The structure was curated by removing the long cytochrome b562 domain used for crystallization and co-crystalized membrane components. The ligand and water molecules and sodium present in the receptor core were maintained (except in the D2.50A system, where the allosterically bound sodium was removed, and the Apo system, where the ligand was removed). Protonation of residues was assigned using the Protonate3D module in MOE. The receptor was aligned to the membrane using the OPM database. The generated complex was then embedded in the POPC bilayer and solvated using TIP3 water using the CHARMM-GUI server. The N and C-terminal ends were patched with ACE and CT3 patches, respectively.

Simulations were carried out similarly to previously published protocols(*50*, *51*) with the ACEMD simulation package(*52*). Ligand parameters were assigned by ParamChem from parameters in the CHARMM General force field(*53*, *54*). Protein and lipid parameters were obtained from the Charmm36M(*55*) and Charmm36(*56*), respectively. The systems were first relaxed for 40 ns under constant pressure and temperature (NPT) with a time step of two fs and gradually decreasing harmonic constraints applied to the protein backbone and ligand. Temperature was maintained at 310 K using the Langevin thermostat and pressure was kept at one bar using the Berendsen barostat. The equilibration run was followed by 100 x 128 ns (for the WT, D2.50A and APO systems) or 50 x 128 ns (for all other systems) production runs under constant volume and temperature (NVT), with a 4 fs time step. Simulation analysis was carried out in the Visual Molecular Dynamics package(*57*) while ligand-receptor interactions were quantified using in-house scripts and the get-contacts package [https://github.com/getcontacts/getcontacts. For the analysis, the initial 64 ns of each replicate were discarded to exclude analysis of potential equilibration events of the system.

For RMSF analysis presented in Figures 1 and 4, the resulting difference between the studied system was normalized with min–max feature scaling, using the following formula:

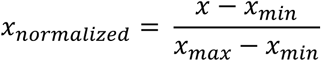

All simulation data is available in the GPCRmd server (https://www.gpcrmd.org/dynadb/publications/1735/).

### Potential of Mean Force

To calculate the unbiased free energy surface associated with conformational changes of Y7.53, we analyzed the simulations with Python Jupyter Notebooks(*58*)(*59*) using PyEMMA. To describe the conformation of Y7.53 we have selected the distances between the OH atom of Y7.53 and Cα atoms of N1.50 and N8.47. The collective variables were discretized into 20 equally spaced bins. We employed the transition-based reweighting analysis method (TRAM)(*60*) with default initialization and a lag time of 1ns. We allowed a maximum number of 25000 self-consistent iterations and the convergence criterion based on the maximal free energy was set to 1E-15 in *k*_B_*T* units.

### Molecular Biology

A codon-optimized DOR-Tango construct (Addgene #66461) was used as a template for mutagenesis generation. Single-point mutations were introduced using the QuikChange Mutagenesis Kit (Agilent, ON), and the resulting mutants were verified through Sanger sequencing.

### Cell Culture

HEK293T cells were cultured in Dulbecco’s Modified Eagle Medium (DMEM) supplemented with 5% fetal bovine serum (FBS), 5% bovine calf serum (BCS), and 100 μg/mL penicillin-streptomycin. The HTLA cells, generously provided by Dr. Richard Axel, are HEK293T cells stably expressing human β-arrestin fused to Tobacco Etch Virus (TEV) protease and a luciferase reporter gene. These cells were cultured in DMEM enriched with 5% FBS, 5% BCS, 100 μg/mL penicillin-streptomycin, 2.5 μg/mL puromycin, and 50 μg/mL hygromycin. All cell lines were maintained at 37 °C in a humidified environment with 5% CO2.

### Tango β-Arrestin Recruitment Assay

The assay was conducted using an adapted version of the Tango protocol previously described in prior studies. HTLA cells were transfected using the PEI precipitation technique. Briefly, 1.5 × 10⁶ cells were seeded in a 6-well plate with 2 mL of growth medium, transfected with 2 μg of DNA mixed in 200 μL Opti-MEM and 6 μL PEI (1 mg/mL, pH 7.0) after a 20-minute incubation. The following day, cells were seeded into Poly-l-Lysine (PLL)-coated 384-well white, clear-bottom plates at a density of 15,000 cells per well, with 40 μL of DMEM containing 1% dialyzed FBS per well. On the third day, ligand solutions were prepared at 3× concentration in filtered assay buffer (20 mM HEPES, 1× HBSS, pH 7.40) and applied to the cells at a volume of 20 μL per well for an overnight incubation lasting 16–20 hours. Media and ligand solutions were subsequently removed, and 20 μL of freshly prepared Glo reagent was added to each well; the reagent composition included 108 mM Tris–HCl, 42 mM Tris-base, 75 mM NaCl, 3 mM MgCl2, 5 mM dithiothreitol (DTT), 0.2 mM coenzyme A, 0.14 mg/mL d-luciferin, 1.1 mM ATP, 0.25% (v/v) Triton X-100, and 2 mM sodium hydrosulfite. Plates were incubated for 10 minutes at ambient temperature before luminescence quantification, which was performed using a Hidex Sense Beta Plus reader (Gamble Technologies, ON). Data were analyzed via nonlinear least-squares regression using the sigmoidal dose-response function in GraphPad Prism 9.0. Results from three independent experiments (n = 3) conducted in quadruplicate are presented as Relative Light Units (RLUs) or normalized values, 128as indicated in the figure legends.

### Analysis of Cell Surface Expression via ELISA

HEK293T cells were seeded into 6-well plates and transfected with WT DOR and mutant GPCR-Tango constructs. Following transfection, the cells were reseeded into 384-well plates at a density of 30,000 cells per well and incubated with or without 10 nM DADLE for 18 hours. The next day, cells were fixed with 20 μL of 4% paraformaldehyde for 10 minutes, followed by blocking with 20 μL of 5% normal goat serum in PBS for 30 minutes. After blocking, an anti-FLAG-HRP conjugated antibody (Sigma), diluted 1:10,000, was applied at 20 μL per well and incubated for 1 hour. The wells were then washed twice with 80 μL of PBS. Luminescence was measured after adding Supersignal ELISA Femto Substrate (Fisher). Error bars indicate the standard deviation (SD) derived from four independent measurements. Data is plotted in Fig. S3

## Author contributions

Conceptualization: TMS, PMG, JS; Data curation: TMS, MZ, IRE, GF; Formal analysis: TMS, MZ, IS, EK, MLB, MMS; Funding acquisition: TMS, PMG, JS; Investigation: TMS, MZ, IS, MMS; Methodology: TMS, MZ, IS, MMS, PMG, JS; Project administration: TMS, PMG, JS; Resources: TMS, PMG, JS; Validation: TMS, MZ; Visualization: TMS, MZ, IS, EK, MLB, Writing original draft: all authors

## Acknowledgments

TMS acknowledges support from the PRELUDE grant 2017/27/N/NZ2/02571 financed by the National Science Center of Poland and Project PID2023-146609OA-I00 financed by MICIU/AEI /10.13039/501100011033 and by FEDER, UE, MZ and PG acknowledge support from the Canadian Institutes of Health Research and the Natural Sciences and Engineering Research Council of Canada.

## Supplementary figures

**Figure S1.**
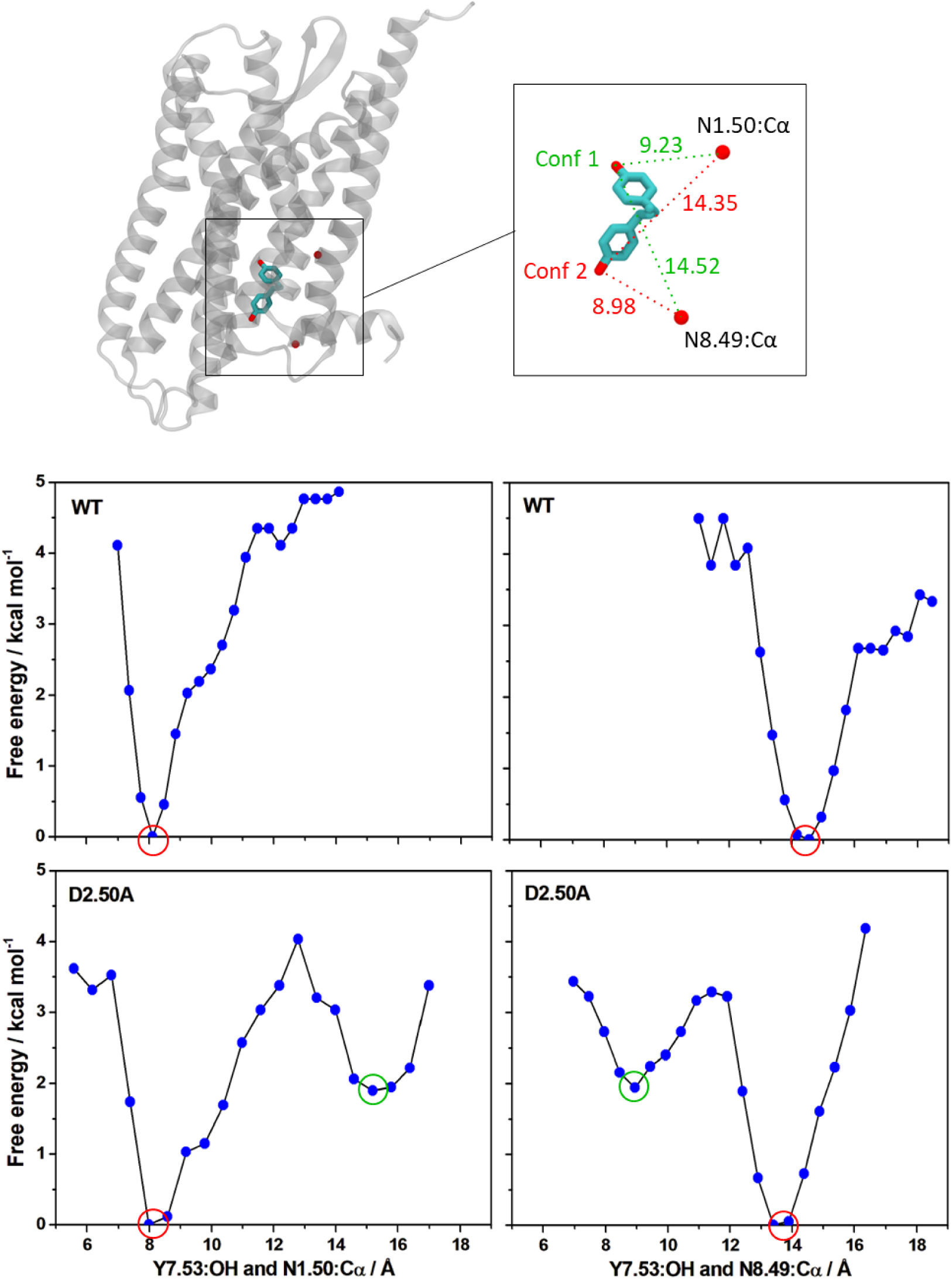
Unbiased free energy surfaces for the conformational rearrangements of Y7.53 obtained from WT and D2.50A simulations. The conformation of Y7.53 is described using two collective variables: the distances between the hydroxyl oxygen of Y7.53 and the Cα atoms of residues N1.50 and N8.47. For comparison, the distance values corresponding to conformation 1 and conformation 2, as defined in this manuscript, are indicated (top).

**Figure S2.**
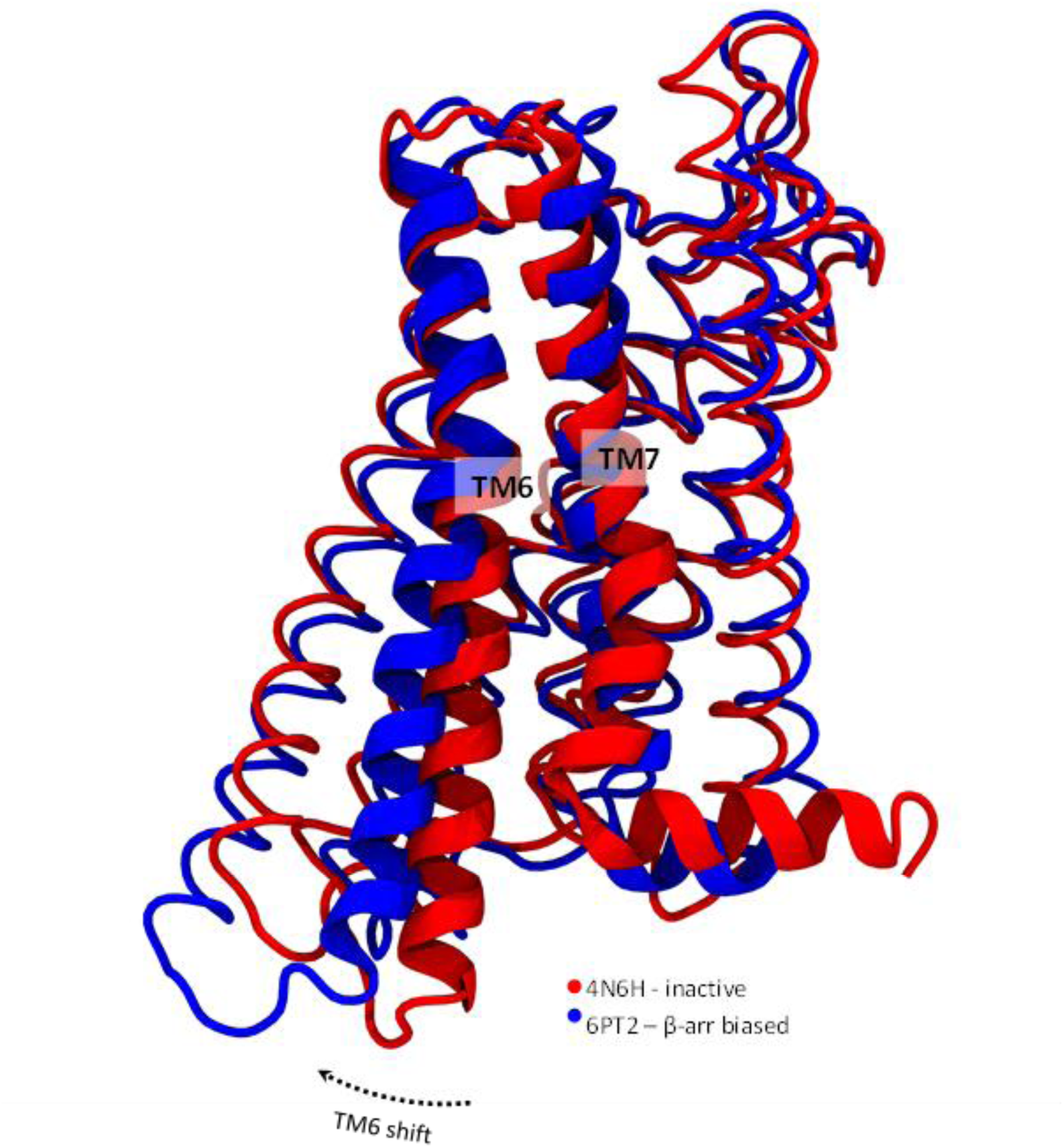
Comparison of TM6 conformation between δOR structures in the inactive (red) and β-arr biased (blue) states. The observed TM6 shift is highlighted with an arrow.

**Figure S3.**
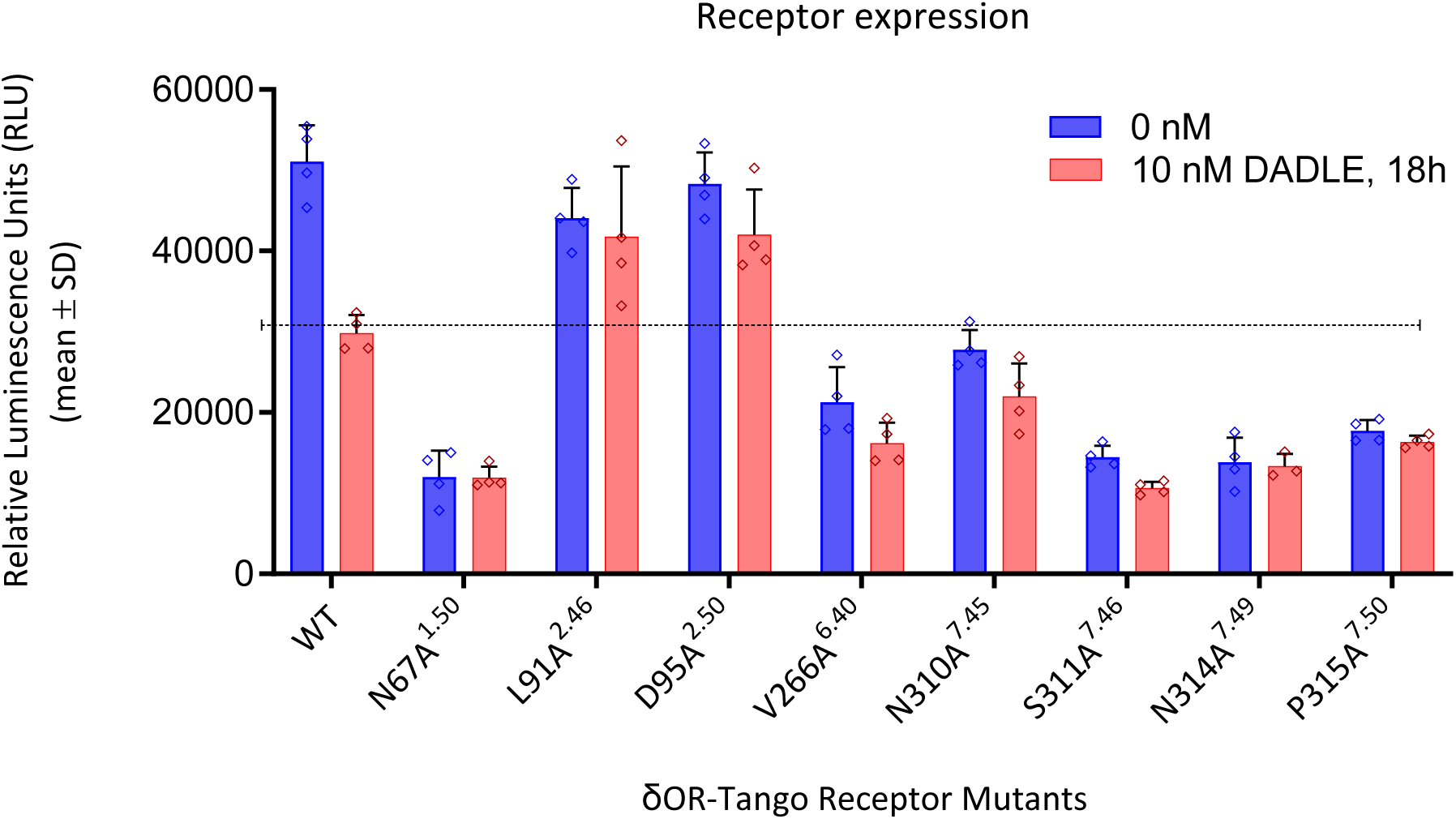
Analysis of Cell Surface Expression via ELISA. Error bars indicate the standard deviation (SD) derived from four independent measurements.

**Table S1.** Summary of simulations carried out in the study.

| System | Number of replicates | Simulation time |
| --- | --- | --- |
| WT + Naltrindole | 100 x 128 ns | 12,8 μs |
| D2.50A + Naltrindole | 100 x 128 ns | 12,8 μs |
| Apo D2.50A | 100 x 128 ns | 12,8 μs |
| N1.50A + Naltrindole | 50 x 128 ns | 6,4 μs |
| L2.46A + Naltrindole | 50 x 128 ns | 6,4 μs |
| V6.27A + Naltrindole | 50 x 128 ns | 6,4 μs |
| N7.45A + Naltrindole | 50 x 128 ns | 6,4 μs |
| S7.46A + Naltrindole | 50 x 128 ns | 6,4 μs |
| N7.49A + Naltrindole | 50 x 128 ns | 6,4 μs |
| P7.50A + Naltrindole | 50 x 128 ns | 6,4 μs |
| Aggregated simulation time: |  | 83,2 μs |

